# Identification of novel inhibitors of *Mycobacterium smegmatis* growth through genome-wide overexpression of Cluster P3 mycobacteriophage Xavia genes

**DOI:** 10.64898/2026.01.06.698002

**Authors:** Anushka Tennakoon, Iresha K. Edirisingha, Ethan Rutledge, Ankita Bhattacharyya, Ella King, Madelyn Moresi, Amara Lovings, Parker R. Yarborough, Anna Hoben, Garrett Manns, Toney Ray Gibson, Syed Mohammed Ashfaque Uddin, Yeseul Bae, Natalie Taylor, Taylor L. Johnson, Madelyn Futral, Paige Wilson, Jalyn Brown, Zackari L. Arbogast, Abigail Brooks, Carson Ward, Courtney Foxworth, Kacey Nguyen, Daphne Fairchild, Deisy Lemus, Hayley White, Holly Craft, Justin Adonis, Kamiya Givan, Sergio Valdivia, Kayci Beth Wallace, Madeline Bent, Dmitri Mavrodi, Danielle M. Heller, Ramesh Rijal

**Affiliations:** School of Biological, Environmental and Earth Sciences, The University of Southern Mississippi, Hattiesburg, MS 39406, USA; Wheat Health, Genetics and Quality Research Unit, U.S. Department of Agriculture, Agricultural Research Service, Pullman, WA 99164, USA; Department of Science Education, Howard Hughes Medical Institute, Chevy Chase, MD 20815, USA

**Keywords:** *mycobacteriophage*, *Mycobacterium smegmatis*, *cytotoxicity*, *genome-wide screen*

## Abstract

We examined whether genes encoded by the mycobacteriophage Xavia disrupt growth of *Mycobacterium smegmatis*, a widely used mycobacterial model. Seventy-one Xavia genes were individually expressed using an inducible plasmid system and assessed for effects on colony formation. Two genes were lethal even without induction, indicating toxicity under basal expression. Induction of sixteen additional genes reduced bacterial growth, spanning structural proteins, lysogeny regulators, DNA-associated enzymes, a lysis protein, and several genes with no known function. These findings expand functional insights into mycobacteriophage gene repertoires and identify candidates for future mechanistic studies.

**Abstract:** Bacteriophage genomes encode large numbers of genes with no known function, and many of these genes affect essential host processes when expressed in a heterologous system. For mycobacteriophages, genome-wide overexpression in *Mycobacterium smegmatis* provides a direct way to identify proteins that impair growth and to determine which mycobacterial pathways are sensitive to phage gene products. To evaluate the cytotoxic potential of the Cluster P3 phage Xavia, a lineage that has not undergone functional screening, we constructed an arrayed pExTra library containing 71 predicted Xavia genes under control of the anhydrotetracycline inducible promoter *pTet*. All constructs were sequence-verified and transformed into *M. smegmatis*, and induction allowed measurement of gene-specific effects on growth. Two genes prevented recovery of transformants, suggesting toxicity under basal promoter leakiness. Inducible expression of 16 additional genes impaired growth, and these inhibitory proteins include structural components, regulators of lysogeny, enzymes of DNA metabolism, a lysis factor, and several proteins with no known function. Four of the strongest inhibitors were genes with no known function. These results extend functional screening into the previously untested P3 branch of Actinobacteriophages and identify new proteins that require mechanistic analysis.

## Introduction

Bacteriophages have co-evolved with bacterial hosts for billions of years, producing highly mosaic genomes shaped by extensive horizontal gene transfer (Pedulla et al., 2003; Russell & Hatfull, 2017). Phage-encoded genes have generated key molecular tools, including restriction enzymes, CRISPR-Cas systems, and regulators of bacterial immunity (Ahmed et al., 2018; Al-Ouqaili et al., 2025; Cong et al., 2013; Hampton et al., 2020; Loenen et al., 2014; Makarova et al., 2011; Mali et al., 2013; Roberts, 2005). Many phage proteins act directly on host pathways, which control envelope integrity, DNA replication, and stress responses (Darwin, 2013; Dunne et al., 2021; Frye et al., 2005; Kim et al., 2024; Payne et al., 2009; Stone et al., 2019). At the same time, most phage genomes encode large numbers of proteins with no recognizable sequence features (Hatfull, 2015, 2020). These “no known function” (NKF) genes represent a reservoir of unexplored activities that may alter host physiology or antimicrobial susceptibility.

To organize the diversity of actinobacteriophage proteins, predicted proteins are grouped into phamilies (phams) based on amino-acid similarity (Cresawn et al., 2011; Gauthier et al., 2022). Previous SEA-GENES overexpression screens showed that cytotoxic no known function genes are often enriched in early lytic regions, which supports roles in host takeover and establishes genome-wide screening as an effective approach for mapping phage gene function (Amaya et al., 2023; D. Heller et al., 2022; D. M. Heller et al., 2024; Ko & Hatfull, 2018; Pollenz et al., 2024; Tafoya et al., 2025).

Mycobacteria include important human pathogens such as *Mycobacterium tuberculosis* and non-tuberculous mycobacteria, many of which show intrinsic antibiotic tolerance and limited treatment options (Boldrin et al., 2020; Goossens et al., 2020; Nguyen, 2016; Wallis et al., 1999). Because direct genetic screening in *M. tuberculosis* is not practical, *M. smegmatis* serves as a well-established surrogate host for identifying conserved vulnerabilities in mycobacterial physiology (Altaf et al., 2010; Lelovic et al., 2020; Sparks et al., 2023; T et al., 2020). Phage genes that impair *M. smegmatis* growth can identify conserved processes that are broadly essential across mycobacteria (Sparks et al., 2023).

Here, we present a genome-wide overexpression analysis of the Cluster P, Subcluster P3 mycobacteriophage Xavia, a temperate siphovirus encoding 71 genes, of which only 39 have predicted functions. We identified phage genes that impair *Mycobacterium smegmatis* growth, placed them in functional and phamily context, and compared their distribution to previously screened mycobacteriophages (Amaya et al., 2023; Heller et al., 2022; Pollenz et al., 2024; Tafoya et al., 2025). Cluster P phages are genetically distinct from previously examined Cluster K and Cluster F phages and have not previously undergone functional cytotoxicity screening. Two Xavia genes were lethal under basal promoter leakiness and could not be recovered as transformants, while inducible overexpression of 16 additional genes, representing approximately 23 percent of the genome, produced measurable cytotoxicity, including seven with severe growth inhibition. This study extends SEA-GENES functional coverage into a previously untested phage cluster and identifies novel phage gene products with cytotoxic activity.

## Materials and methods

### Growth of mycobacteria and mycobacteriophage

*Mycobacterium smegmatis* mc^2^155 was grown at 37 °C in Middlebrook 7H9 (Difco) broth supplemented with 10% AD (2% w/v dextrose (Fisher Chemical), 145 mM NaCl (Fisher Chemical), 5% w/v albumin fraction V (Millipore), 0.05% Tween 80 (Fisher Scientific), 10 µg/mL cycloheximide (Acros Organics), and 50 µg/mL carbenicillin (Research Products International) or on Middlebrook 7H11 (Difco) agar supplemented with 10% AD, 0.5% glycerol (Fisher Scientific), 10 µg/mL cycloheximide, and 50 µg/mL carbenicillin. The competent cells were prepared according to the SEA-GENES Instructor Guide (https://seagenesinstructorguide). A single isolated colony of *M. smegmatis* mc^2^155 was grown overnight (∼ 16 h) in 5 mL of 7H9 complete medium at 220 rpm and 37 °C until the cultures reached an OD between 1 and 3. Then, 400 mL of 7H9 complete with Tween 80 (0.05 %) was inoculated with the *M. smegmatis* mc^2^155 overnight culture, at a starting optical density (OD) of 0.005. The new culture was allowed to incubate at 37 °C while shaking at 220 rpm overnight for 16-24 hours until the culture reached an OD of ∼ 0.6. Then, the cells were incubated on ice for 1.5 hours. The chilled cultures were then centrifuged at 2,000 x g for 20 minutes at 4 °C. The supernatant was discarded, and the cells were washed with 200 mL of ice cold 10 % glycerol three times at 2,000 x g for 20 minutes at 4°C. After the final wash, the cells were resuspended in 1:250 of the original volume with ice-cold 10 % glycerol and used in transformation.

To transform cells with pExTra plasmids (Heller et al., 2022), 100 ng of pExTra-Xavia gene plasmid DNA (described below) was mixed with 50 µL of electrocompetent *M. smegmatis* mc^2^155 cells and electroporated in 0.1 cm cuvettes (Lightlabs) using a Bio-Rad Laboratories GenePulser Xcell electroporator (1.8 kV, 10 µF, 600 Ω) (Bio-Rad). Following transformation, cells were resuspended in 1 mL of 7H9 medium and incubated at 37 °C for 1 h. The culture was spun down at 2000 x g for 5 minutes and 900 µL of the supernatant was discarded. The pelleted transformed cells were then resuspended in the remaining 100 µl of the media and then plated on 7H11 agar supplemented with 5 µg/mL kanamycin. Plates were incubated at 37 °C for up to 5 days, and colony growth was monitored throughout the incubation period.

### Construction of the pExTra Xavia library

Individual Xavia genes were inserted into the pExTra shuttle plasmid under control of the anhydrotetracycline (aTc) responsive *pTet* promoter, with the transcriptional reporter gene *mCherry* positioned immediately downstream of the cloned insert (**Fig. 1a**; Heller et al., 2022). Xavia genes were PCR-amplified using gene-specific primers (Integrated DNA Technologies) (Supplementary Table 1), Q5 DNA polymerase (New England Biolabs Q5 HotStart 2× Master Mix), and a high-titer Xavia lysate. Gene-specific primers anneal to the first and last 15–25 bp of each gene sequence and introduce a uniform ATG start codon (forward primers) or TGA stop codon (reverse primers). Each primer was also designed to contain regions of identity to the pExTra01 plasmid; forward primers contain a uniform RBS and 5′ 21 bp of homology to pExTra01 downstream of the *pTet* promoter, and all reverse primers contain a separate 5′ 25 bp of homology to pExTra01 upstream of *mCherry* (Supplementary Table 1). Homologous sequences were used to direct insertion of Xavia gene fragments into the pExTra plasmid downstream of the *pTet* promoter by isothermal assembly with the NEB HiFi 2× Master Mix (Heller et al., 2022). Linearized pExTra01 plasmid was prepared for isothermal assembly reactions via PCR (NEB Q5 HotStart 2× Master Mix) of pExTra01 using divergent primers pExTra_F and pExTra_R. Recombinant plasmids were recovered by transformation of *Escherichia coli* NEB5α F′I^Q^ (New England Biolabs) and selected on LB agar supplemented with 5 µg/mL kanamycin. Sequence integrity of all pExTra–Xavia plasmids was confirmed by Sanger sequencing at Azenta using pExTra_universalR and pExTra_seqF primers, and longer inserts were further validated with internal primers provided in Supplementary Table 1. All plasmid inserts matched the published genome sequence.

**Figure 1.**
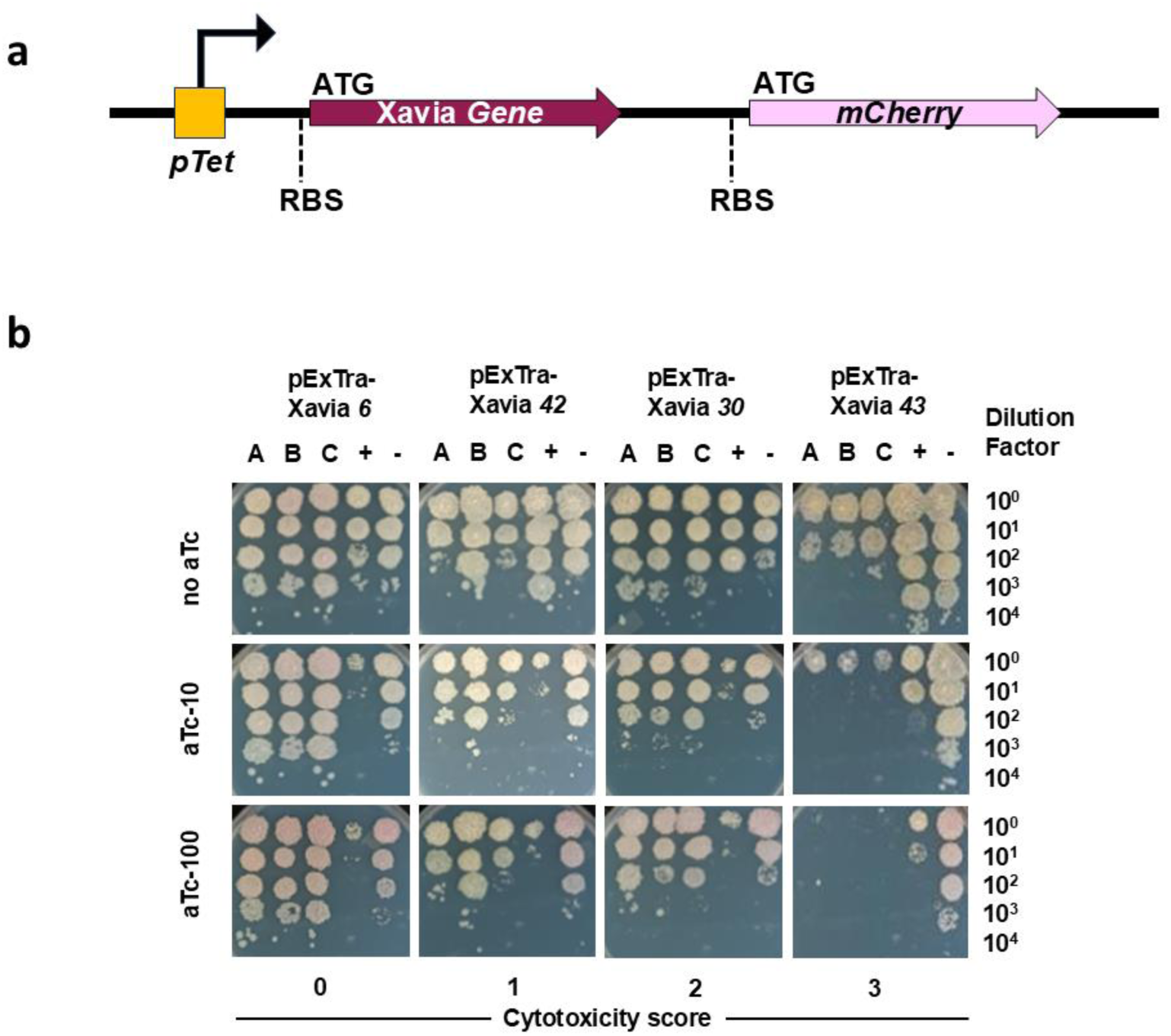
Representative cytotoxicity phenotypes showing Xavia genes with scores 0-3. (a) pExTra constructs used in this study contain each Xavia gene downstream of the aTc-inducible *pTet* promoter and upstream of *mCherry* in a transcriptionally linked operon. Each open reading frame contains its own ribosome-binding site (RBS) and translation start and stop codons. (b) Representative cytotoxicity assays illustrate the range of observed phenotypes. For each gene, three independent transformants (A, B, C) were grown, serially diluted, and spotted on 7H11 agar containing 0, 10, or 100 ng/mL aTc. Positive control strains carried pExTra02 (Fruitloop 52), and negative controls carried pExTra03 (Fruitloop 52-I70S). Cytotoxicity score 0 is represented by Xavia 6, score 1 by Xavia *42*, score 2 by Xavia *30*, and score 3 by Xavia *43*. Images shown are representative of at least two independent experiments.

### Cytotoxicity screen and phenotype scoring

Three independent transformed colonies were selected per each gene and serial dilutions (10⁰–10⁻⁵) cultures were prepared in 7H9, and 10 µL of each dilution was spotted onto 7H11 plates supplemented with Kanamycin and 0, 10, or 100 ng/mL aTc, as described in previous SEA-GENES studies (Amaya et al., 2023; Heller et al., 2022; Pollenz et al., 2024; Tafoya et al., 2025). pExTra-Fruitloop*52* wild type (toxic positive control) and the mutant (I70S; non-toxic negative control) were spot plated in serial dilutions along with the three replicates (Heller et al., 2022; Ko & Hatfull, 2018). The growth of *M. smegmatis* was observed over five days. Cytotoxic phenotypes were categorized using the scoring framework established in earlier SEA-GENES studies of Amelie, Hammy, Waterfoul, and Girr (Amaya et al., 2023; Heller et al., 2022; Pollenz et al., 2024; Tafoya et al., 2025). Strains showing growth comparable to the negative control were assigned a score of 0. A visible reduction in colony size was scored as 1. A decrease of roughly one to two orders of magnitude in colony number was scored as 2. A loss of three logs or more in recoverable colonies, or the near absence of growth, was scored as 3. For constructs assigned score 0, the presence of pink colonies on aTc plates confirmed induction of the *pTet* promoter and successful *mCherry* expression. We performed 2-4 independent experiments to validate the observed cytotoxicity. When variability occurred, particularly among weak phenotypes, the mildest reproducible effect was used as the final score.

Pink coloration arises from transcription of the downstream *mCherry* reporter and reflects promoter induction by aTc. Thus, pink colonies on aTc plates confirm successful transcriptional activation of the *pTet*–Xavia gene–*mCherry* cassette, even when no cytotoxic phenotype is observed. Occasional weak *mCherry* fluorescence was observed on plates without aTc due to basal promoter leakiness, a phenomenon reported in Waterfoul, Hammy, and Girr screens (Amaya et al., 2023; Heller et al., 2022; Pollenz et al., 2024). This leakiness was not sufficient to affect scoring but confirmed promoter activity.

### Xavia genomic analysis

The Xavia genome map was generated using Phamerator (Cresawn et al., 2011), which displays coding regions, functional annotations, and shared phamilies with related phages. Gene functions were taken from the published Xavia GenBank record (Accession MH230879), and no additional annotation refinement was required beyond the existing database entries. As a P3 cluster phage, Xavia contains few genes with close homologs in other SEA-GENES phages (Amaya et al., 2023; Heller et al., 2022; Pollenz et al., 2024; Tafoya et al., 2025), and only a limited number of phamilies are shared with Girr, Amelie, Waterfoul, or Hammy (**Table 1**).

**Table 1.** Cytotoxic Xavia genes identified by heterologous expression in *M. smegmatis*.

| Xavia gene | Phamily<sup>a</sup> | Length (aa) | Predicted function and features | Cytotoxicity | Homologs in other phages<sup>c</sup> |
| --- | --- | --- | --- | --- | --- |
| **4** | 665 | 446 | Head maturation protease | 1 |  |
| **5** | 258736 | 121 | Scaffolding protein | 1 |  |
| **29** | 261076 | 120 | Minor tail protein | 1 | Girr gp35 (Pollenz et al., 2024) |
| **42** | 260356 | 265 | Antirepressor | 1 |  |
| **3** | 130463 | 486 | Portal protein | 2 |  |
| **18** | 59 | 581 | Minor tail protein | 2 | Girr gp15 (Pollenz et al., 2024) |
| **30** | 14397 | 375 | DNA methylase | 2 |  |
| **47** | 261119 | 119 | NKF | 2 |  |
| **55** | 253641 | 143 | RusA-like resolvase | 2 |  |
| **27** | 261033 | 388 | Lysin B | 3 |  |
| **31** | 14469 | 321 | NKF | 3 |  |
| **37** | 110332 | 144 | NKF | 3 |  |
| **39** | 259157 | 192 | Immunity repressor | 3 |  |
| **43** | 261346 | 66 | Excise | 3 |  |
| **48** | 115328 | 69 | NKF | 3 |  |
| **51** | 106454 | 118 | NKF | 3 |  |
| **58** | 85836 | 131 | NKF | 3 |  |
| **62** | 200918 | 297 | NKF | 3 |  |

Cytotoxicity was measured by comparing growth of each pExTra–Xavia strain on 7H11 agar containing 0, 10, or 100 ng/mL anhydrotetracycline. Scores reflect reproducible reductions in colony size or viability relative to the same strain plated without inducer. Functional annotations and phamily assignments were obtained from Phamerator and PhagesDB. NKF indicates a gene product with no known function. <sup>a</sup>Phamily numbers were obtained from PhagesDB (accessed 28 September 2025). <sup>b</sup>Cytotoxicity scores reflect reproducible growth inhibition at 10 or 100 ng/mL aTc: 0, no effect; 1, reduced colony size; 2, ~1–2 log reduction; 3, ≥3-log reduction. <sup>c</sup>Homologs shown correspond to genes in previously screened SEA-GENES phages.

Gene content similarity between Xavia and these phages was quantified using the PhagesDB gene content tool (https://phagesdb.org/genecontent/). Xavia exhibited low gene content similarity with Girr (13.14%), Amelie (6.77%), Waterfoul (7.42%), and Hammy (4.94%), consistent with its placement in a distinct cluster. Phamily assignments were downloaded from the PhagesDB database on 28 September 2025 and used to compare the distribution of shared and unique gene groups.

Functional predictions for Xavia genes were carried out using the same workflow applied in previous SEA-GENES studies (Amaya et al., 2023; Heller et al., 2022; Pollenz et al., 2024; Tafoya et al., 2025). Gene neighborhoods, phamily assignments, and genome maps were examined using Phamerator (Cresawn et al., 2011). Pairwise sequence comparisons were performed using BLASTp (NCBI), and hits with E-values < 1×10⁻⁵, >50% coverage, and strong percent identity were considered meaningful. Domain architecture was evaluated using HHpred (Gabler et al., 2020) with the PDB_mmCIF70_25_May database. Predicted protein structures were generated using AlphaFold3 (alphafoldserver.com), and structural similarity searches were performed in Foldseek (van Kempen et al., 2024). Proteins lacking confident matches in PDB database were classified as hypothetical proteins/NKFs. Bioinformatic results used for functional grouping of cytotoxic genes are summarized in Supplementary Table 2.

Xavia showed no identifiable clustering of cytotoxic or functional modules within the genome map. Cytotoxic genes were dispersed throughout structural, lysogeny, DNA metabolism, and NKF regions. This dispersed organization is consistent with the high genomic diversity typical of P-cluster phages (Russell & Hatfull, 2017) and supports the observation that Xavia shares only a small subset of phamilies with the other phages evaluated in SEA-GENES studies (Amaya et al., 2023; Heller et al., 2022; Pollenz et al., 2024; Tafoya et al., 2025).

## Results and discussion

### Inducible expression of Xavia genes identifies strongly cytotoxic and lethal activities in *M. smegmatis*

To investigate the cytotoxic effects of bacteriophage Xavia gene on growth of *M. smegmatis*, a total of 71 genes were individually cloned into the pExTra shuttle plasmid (Heller et al., 2022), which contains a *pTet*-inducible promoter and a linked *mCherry* transcriptional reporter gene (**Fig. 1a**). The resulting recombinant constructs were selected based on kanamycin resistance and verified by sequencing. All inserts were confirmed in *E. coli*, ruling out cloning artifacts before transformation. The verified plasmids were subsequently transformed into *M. smegmatis*. However, two genes, Xavia *37* and Xavia *48*, could not be successfully transformed despite repeated attempts, indicating that basal promoter leakiness in the *pTet* system (Amaya et al., 2023) is likely sufficient to prevent growth of the transformants. Bioinformatic analysis indicates that gp37 shares limited sequence similarity with a protein annotated as a toxin in *Prescottella equi*, with approximately 38 percent amino acid identity, as summarized in Supplementary Table 2. Expression of gp37 was sufficient to prevent recovery of transformants, which is consistent with a strong inhibitory effect on bacterial growth (Freeman et al., 2024). In contrast, recovery of stable transformants may be observed when gp37 is co-expressed with gp38, a gene annotated in PhagesDB as a putative antitoxin. By comparison, gp48 lacks an associated antitoxin and shows the same severe toxicity reported for other NKF genes such as Waterfoul gene 8 (Heller et al., 2022) and Hammy gene 9 (Amaya et al., 2023).

For the remaining 69 genes, three independent transformed colonies per gene were suspended in 7H9 broth, serially diluted, and spotted onto 7H11 agar supplemented with 0, 10, or 100 ng/ml aTc. Cytotoxicity scores ranged from 0 to 3 based on colony size and log-fold reductions in viability. Strains expressing the pExTra plasmid carrying the cytotoxic Fruitloop *52* gene (Ko & Hatfull, 2018) served as the positive control, while strains expressing the pExTra plasmid encoding the non-toxic Fruitloop *52*-I70S (Heller et al., 2022) were used as the negative control, and both controls behaved consistently across experiments.

All cytotoxicity scores were calculated by comparing viability of strains on 100 ng/ml aTc relative to the no-aTc control, following the same scoring framework used in previous Waterfoul, Hammy, and Girr screens (Amaya et al., 2023; Heller et al., 2022; Pollenz et al., 2024). Representative genes are shown in **Figure 1b**. Induction with 10 or 100 ng/ml aTc did not alter colony size or numbers for Xavia *6* (score = 0), although minor natural variation was observed among replicates. For Xavia *42*, induction with 10 or 100 ng/ml aTc reduced colony size (score = 1), and this reduction was consistent across plates. Induction of Xavia *30* with 10 or 100 ng/ml aTc reduced colony numbers by approximately 1-log and 2-log, respectively (score = 2). Induction of Xavia *43* with 10 or 100 ng/ml aTc caused ≥ 3-log reductions in colony numbers (score = 3). Some constructs showed minor variation in colony size or *mCherry* intensity across independent replicates, a pattern also noted in other SEA-GENES screens (Amaya et al., 2023; Heller et al., 2022; Pollenz et al., 2024; Tafoya et al., 2025). These differences did not alter the overall cytotoxicity assignments, and phenotypes were stable enough to support the reported scoring.

### Inducible expression of Xavia genes identifies cytotoxic and non-cytotoxic activities in *M. smegmatis*

The cytotoxicity screening identified 18 Xavia genes that inhibited *M. smegmatis* growth to different degrees, and these inhibitory effects occurred across several functional groups. Fourteen of the 18 produced moderate to severe reductions in colony size or number (**Fig. 2**, **Table 1**). The proportion of cytotoxic genes in Xavia was 23%, which is similar to frequencies reported in Hammy (25%) (Amaya et al., 2023), Girr (28%) (Pollenz et al., 2024), Amelie (34%) (Tafoya et al., 2025), and Waterfoul (34%) (Heller et al., 2022).

**Figure 2.**
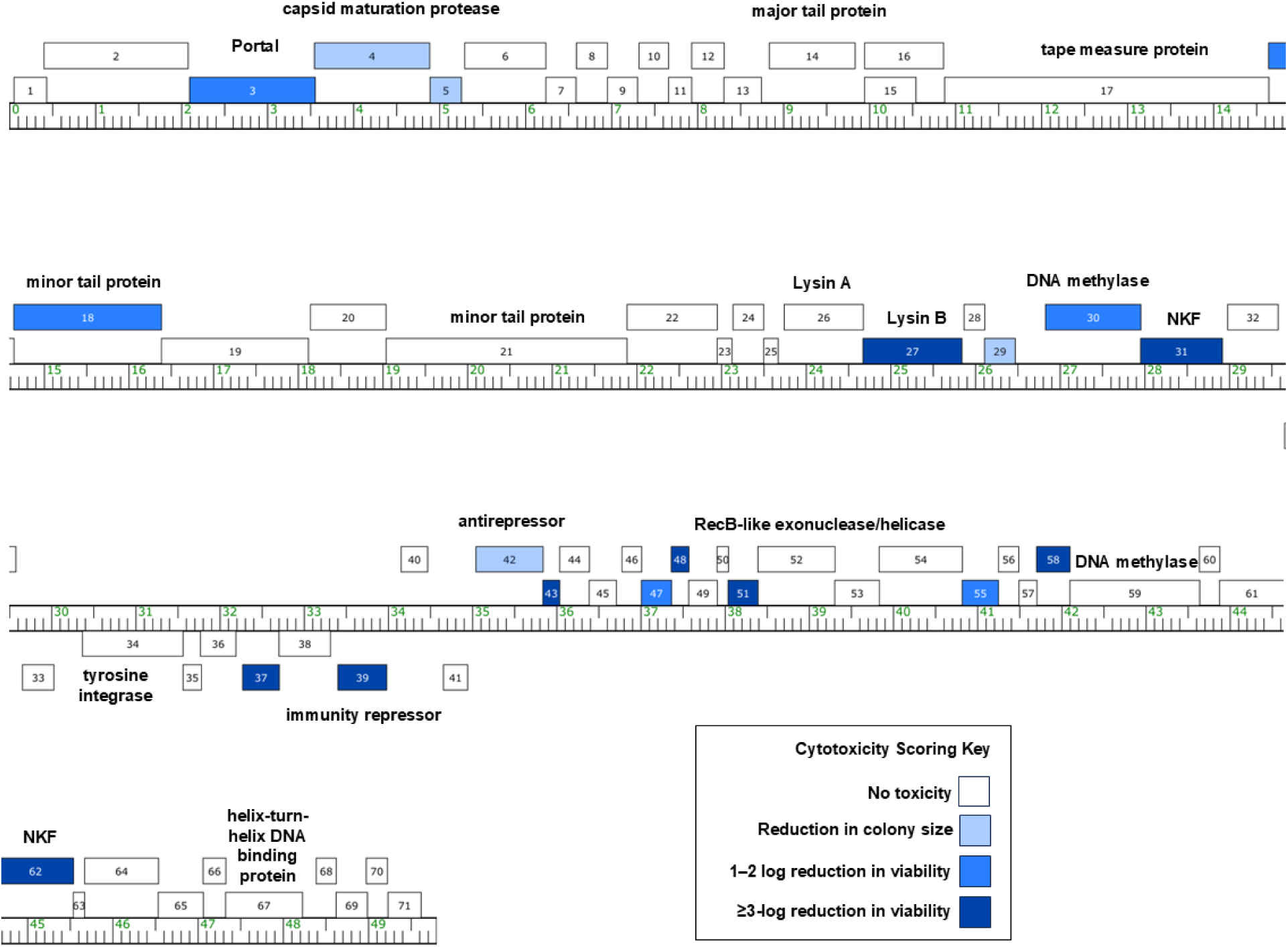
Annotated genome map of mycobacteriophage Xavia with cytotoxicity scores. The Xavia genome is shown with kilobase markers, and each predicted gene is represented by a box, with genes above the line transcribed rightward and those below the line transcribed leftward. Numbers inside each box correspond to gene numbers, and functional assignments follow the Xavia GenBank record. The map was generated using Phamerator. Box shading indicates cytotoxicity scoring, with white boxes designating genes that did not alter *Mycobacterium smegmatis* growth (score 0). Increasing shades of blue indicate greater inhibition: light blue (score 1; reduced colony size), medium blue (score 2; 1–2 log reduction in viability), and dark blue (score 3; ≥ 3-log reduction in viability). A scoring key is provided in the figure.

The 18 identified cytotoxic gene products belonged to multiple functional categories. Five were annotated as structure or assembly proteins, including capsid maturation protease gp4, scaffolding protein gp5, and minor tail proteins gp18 and gp29. Two proteins, DNA methylase gp30 and RusA-like resolvase gp55, have putative roles in DNA replication and metabolism. Several other products have predicted roles in the lysis/lysogeny life cycle decision, including antirepressor gp42, excise protein gp43, immunity repressor gp39, and Lysin B protein gp27. (Pollenz et al., 2024)

Bioinformatic analyses were used to refine the predicted functions of the cytotoxic proteins (Supplementary Table 2). For genes with existing annotations in Phamerator and PhagesDB, BLASTp, HHpred, and structure-based searches supported the assigned roles in structural assembly, lysogeny, and DNA metabolism. Among the NKF proteins, gp31, gp58, and gp62 showed weak but detectable similarity to predicted folds associated with membrane-associated or small enzymatic domains, suggesting potential roles that are not captured by current annotations. In contrast, gp47 and gp51 lacked confident matches in any database and remain completely hypothetical. These NKF genes represent previously unrecognized sources of cytotoxicity, as none of them have reported toxic effects in related SEA-GENES phages. These further analyses did not change GenBank annotations but helped group the cytotoxic genes into functional categories and highlight NKF proteins that may warrant deeper mechanistic studies.

The remaining 53 genes (75%) of the genes tested did not produce cytotoxicity under the overexpression conditions used in this assay, as shown in **Fig. 2** and Supplementary Fig. 1. Most of the non-cytotoxic genes, 44 out of 53, developed pink colonies after aTc induction, which indicates that the gene operon was expressed. Pink coloration reflects promoter activity but does not ensure that the proteins accumulated to levels required for functional effects. The absence of cytotoxicity in these strains may reflect limited protein accumulation, poor solubility of the expressed proteins, or insufficient activity of the expressed proteins, including cases in which cytotoxic functions require combinations of phage proteins rather than single proteins under the growth conditions used in this screen (Cahill & Young, 2019; Mohanraj et al., 2019). Additional validation at the protein level will be needed to determine whether these genes encode stable proteins or whether their products fail to reach levels that affect growth. A small number of constructs showed weak or inconsistent phenotypes that did not meet the criteria for cytotoxicity, and these borderline patterns were also observed in earlier SEA-GENES studies (Amaya et al., 2023; Heller et al., 2022; Pollenz et al., 2024; Tafoya et al., 2025).

### Comparative analysis shows limited phamily conservation but shared cytotoxic functions among mycobacteriophages

To assess whether the cytotoxic genes identified in Xavia share phenotypic similarities with related phages, we compared the corresponding phamilies encoded in Hammy, Waterfoul, Girr, and Amelie (phagesdb.org). Xavia shares 13.14%, 6.77%, 7.42%, and 4.94% gene content similarity with Girr, Amelie, Waterfoul, and Hammy, respectively, resulting in a limited number of shared phamilies (phagesdb.org). Only Girr encoded phams related to cytotoxic proteins identified in Xavia, but the corresponding phamily members in Girr, minor tail protein gp15 and NKF protein gp35 did not produce cytotoxicity (Pollenz et al., 2024). Further investigation is needed to determine whether this phenotypic difference is a consequence of true functional divergence.

Given the low level of gene content similarity between Xavia and previously screened genomes, we looked more generally for consistent functional themes. The scaffolding proteins Girr gp5 and Amelie gp8 (Pollenz et al., 2024; Tafoya et al., 2025) produced similar cytotoxic effects as did minor tail proteins Girr gp19, Amelie gp20, and Waterfoul gp21 (Heller et al., 2022; Pollenz et al., 2024; Tafoya et al., 2025). Conserved roles in DNA metabolism also show up, with DNA methylase Girr gp64 (Pollenz et al., 2024) and RusA-like resolvase Amelie gp58 (Tafoya et al., 2025) exhibiting toxicity. Toxicity of the Xavia immunity repressor gp39 and antirepressor gp42 match observations from Girr showing that its immunity repressor, gp46, and antirepressor, gp48, both of which belong to different gene phamilies than the Xavia products, are highly cytotoxic when overproduced (Pollenz et al., 2024). Immunity repressors from Cluster K phages Hammy, Waterfoul, and Amelie do not cause cytotoxicity, indicating that this phenotype is specific to certain phamilies and not a general consequence of immunity repressor function.

There were some notable differences between Xavia and the other datasets as well. For example, the major capsid proteins from all Cluster K and Cluster F phages tested to date have exhibited toxicity, whereas Xavia gp6 was not observed to inhibit host growth. Furthermore, although multiple lysin A genes have been reported to be toxic when overexpressed individually (Amaya et al 2023, Pollenz et al 2024), to the best of our knowledge, this is the first report of lysin B overexpression impairing host physiology. Altogether, the current screen identified 18 phamilies that impair bacterial growth when overexpressed, and some of these phamilies appear to be novel in the context of cytotoxicity in *M. smegmatis*.

This genome-wide overexpression screen of the P3 cluster mycobacteriophage Xavia provides the first functional characterization reported for a cluster P phage. Despite the low overlap with previously tested gene phamilies in Cluster K and Cluster F phages, the proportion of cytotoxic genes in Xavia, approximately 26% is consistent with frequencies, ranging from 26-34%, reported in Girr (Pollenz et al., 2024), Hammy (Amaya et al., 2023), Amelie (Tafoya et al., 2025), and Waterfoul (Heller et al., 2022). Xavia does not show obvious clustering of cytotoxic genes within the genome, and toxic genes were dispersed among regions associated with structural, DNA metabolism, lysogeny, and NKF functions. These observations support the idea that diverse phage genomes encode a substantial number of proteins capable of impairing host growth when overexpressed, even when their functions are poorly understood. The consistent patterns across independent experiments and agreement with previously reported SEA-GENES datasets support the robustness of the findings. Structural and domain predictions for NKF genes (Supplementary Table 2) suggest that at least some of these cytotoxic proteins adopt folds associated with known cellular functions, even though their specific activities in mycobacteria are still unclear. The relatively high number of cytotoxic genes identified in this study suggests that Xavia uses several parallel mechanisms to impair host physiology. Such functional redundancy is consistent with the diverse host-modulating proteins described in other mycobacteriophages, which frequently encode multiple small genes that independently disrupt membrane integrity, DNA metabolism, or essential growth pathways (Amaya et al., 2023; Freeman et al., 2024; Hatfull, 2018; Heller et al., 2022; M. Iyer et al., 2021; Pollenz et al., 2024; Sharma & Jain, 2025; Tafoya et al., 2025).

## Data Availability

Plasmids generated in this study and their corresponding sequence information can be obtained from the authors upon reasonable request. All data required to evaluate and support the conclusions are provided within the main text, figures, and tables. Additional supporting datasets, including plasmid Sanger sequencing results and validation cytotoxicity assays, are deposited in the SEA-GENES open-access repository GenesDB (https://genesdb.org). These data are accessible through GenesDB after free user registration and can be located by navigating from the homepage to the appropriate phage entry. Supplementary material associated with this article is available through the journal website.

## Acknowledgments

This study was carried out within the framework of the Science Education Alliance GENES program supported by the Howard Hughes Medical Institute. We thank members of the Science Education Alliance community for their contributions. We also thank the University of Southern Mississippi Spring 2024 SEA-GENES cohort (Supplementary Table 3) for conducting cytotoxicity assays to validate data. High-titer Xavia lysates used in this study were provided through the Science Education Alliance and were originally isolated by Stephon Scott at Morehouse College in Atlanta, Georgia. Laboratory reagents and instructional materials for the Microbial Genetics course BSC 477/577L were provided by the University of Southern Mississippi. Additional reagent support was provided by New England Biolabs and Integrated DNA Technologies.

